# Digital Cousins: Simultaneous Optimization of One Model for BMP Signaling in Distant Relatives Reveals Essential Core

**DOI:** 10.1101/2025.02.25.640248

**Authors:** Linlin Li, Thembi Mdluli, Gregery Buzzard, David Umulis

## Abstract

Spatially distributed, nonuniform morphogen gradients play a crucial role in tissue organization during development across the animal kingdom. The Bone Morphogenetic Protein (BMP) pathway, a well-studied morphogen involved in dorsal-ventral (D-V) axis patterning, has been extensively studied in zebrafish, *Drosophila*, and other organisms. Given that this pathway is highly conserved in both form and function, we sought to determine whether a core mathematical model that constrained topology and biophysical parameters could fully reproduce the observed dynamics of gradient formation in both *Drosophila* and zebrafish through changes in expression only. We used multi-objective optimization to simultaneously fit a single core model to *Drosophila* and zebrafish data and conditions. By exploring a single model with varied parameters, we identified both the homology and diversification of the BMP pathway. We find that a core model with only two parametric changes could simultaneously replicate the experimentally measured BMP gradients in both species. This approach, involving simulation and multispecies optimization, provides a rigorous method to identify the minimum parameter adjustments needed for the measurement and simulation of one species to have predictive power in another system. The process offers a framework for enhancing cross-species predictions and improving the utility of preclinical animal models.

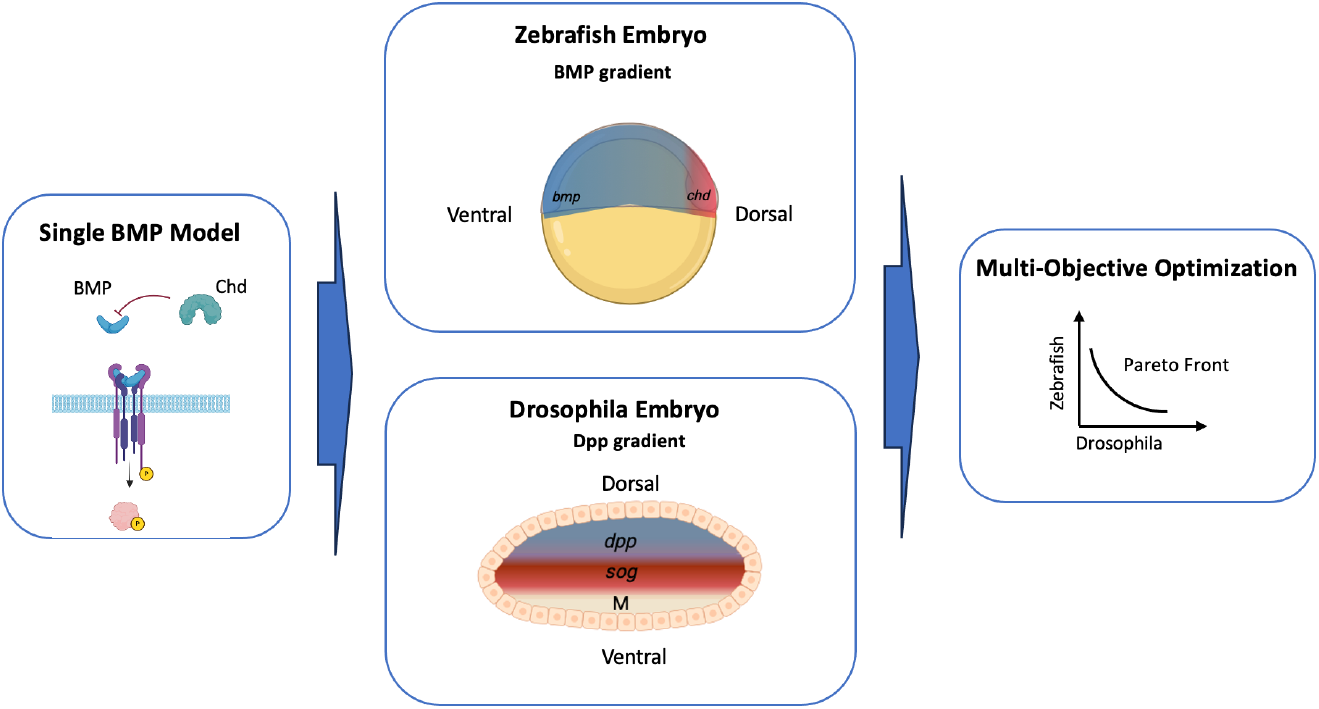

## 1 Introduction

Structures and body plans in organisms across the animal kingdom are specified by spatially distributed nonuniform gradients of molecules called morphogens. These gradients guide the development of patterns from a single-cell structure into a well-organized, patterned organism with diverse structures^1–6^. The precise positioning of patterned features of an organism is crucial, as they emerge from an initially homogeneous group of cells that differentiate in response to morphogen cues. An extensively studied morphogen is the Bone Morphogenetic Protein (BMP), a member of the TGF-β superfamily, which plays a key role in both vertebrates and invertebrates within the animal kingdom. Of current interest is comprehending how BMPs orchestrate the dorsal-ventral (D-V) axis in the early embryos of zebrafish (a vertebrate) and *Drosophila* (an invertebrate). Despite their evolutionary divergence, D-V patterning is established by a common pathway in these diverse species through the interaction of BMP ligands (Decapentaplegic (Dpp)/BMP2/4) with extracellular regulators, especially the antagonistic interaction between regulators of the BMP pathway. Figure 1 illustrates the BMP gradient in patterning the D-V axis in drosophila and zebrafish embryos. The pathway is initiated by a BMP ligand binding and forming a complex with trans-membrane serine-threonine kinase (Type I and Type II) receptors ^7,8^. The binding of BMP ligands to receptors is highly regulated by extracellular factors that both antagonize, Sog/Chordin (Chd), and/or actuate (Tolloid) the signal (Figure 1I). As shown in Figure 1I, upon BMP ligand binding, the Type II receptors phosphorylate the Type I receptors, which in turn phosphorylate members of the R-Smad ^9,10^, including Smad 1, Smad 3, and Smad 8. Phosphorylated R-Smads then bind to Smad 4 to form complexes that translocate to the nucleus, where they accumulate and regulate gene expression ^11–18,^ which leads to precise pattern formation. A simplified representation of the key BMP regulators involved in D-V axis patterning in arthropods and vertebrates is illustrated in Figure 1I.

**Figure 1.**
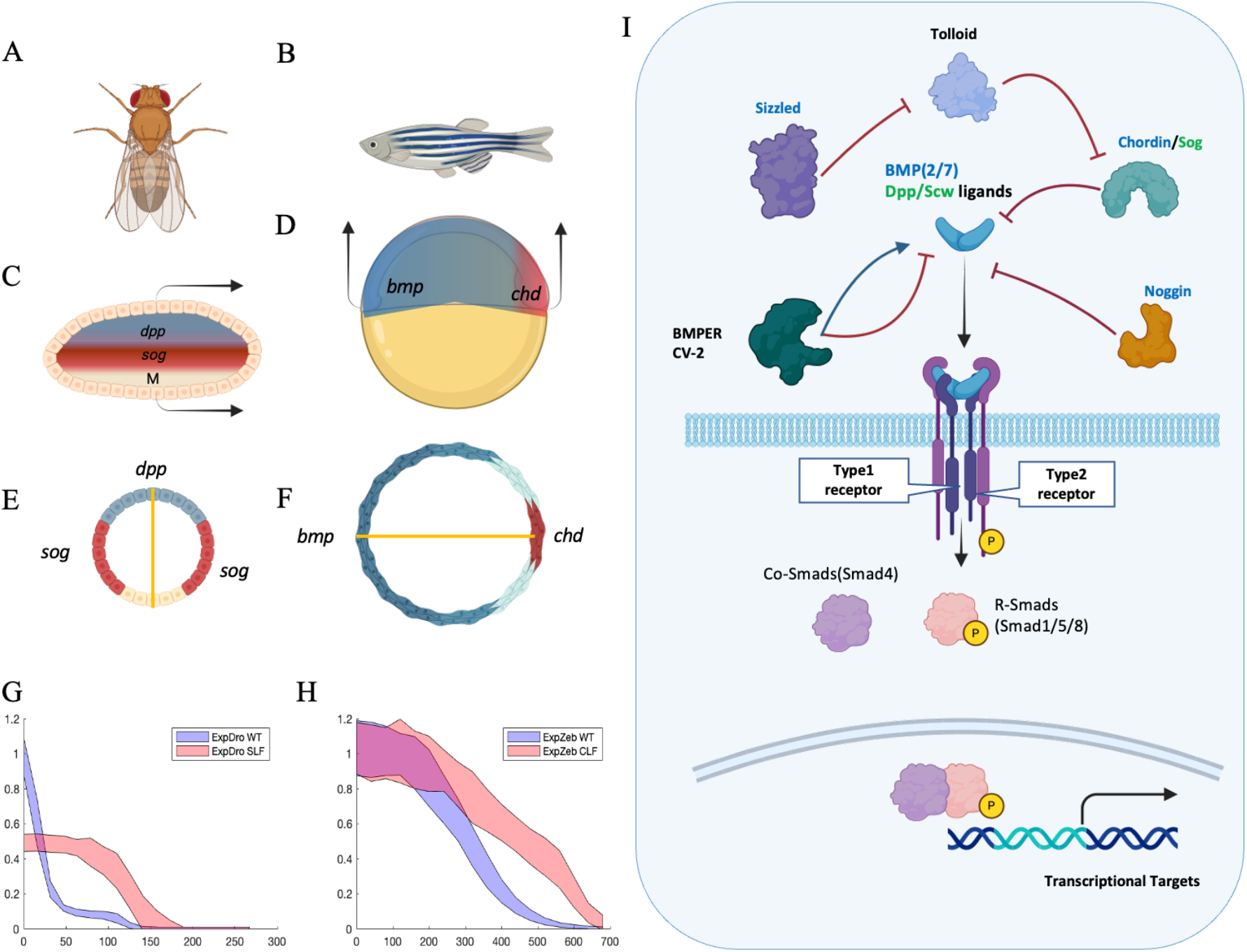
(A) and (B) depict the dorsal-ventral patterning in Drosophila and zebrafish, respectively. (Copyright: National Museum of Natural History), (C) and (D) show the signaling gradient profiles and expression domains of Dpp patterned Drosophila embryo, along with its sectional view (E), and BMP patterned zebrafish embryo and its sectional view (F), (G) presents experimental average profile of P-Smad domains in the Drosophila wild-type embryo(blue) and Sog loss-of-function embryo (red) (275 μm; transverse section). Similarly, (H) shows the experimental average profile of P-Smad domains in the Zebrafish wild-type embryo(blue) and Chordin loss-of-function embryo (red) (700 μm; transverse section). (I) provides a schematic representation of the BMP Smad pathway at the cellular level, highlighting key interactions involved in D-V patterning.

The classical BMP signaling pathway (Figure 1I) demonstrates a conservation of the pathway at the cellular level even though distinct patterns are observed between *Drosophila* and zebrafish at the organism level (Figure G and H). Understanding the mechanism of BMP regulation will elucidate the parts of the signaling pathway that are conserved and those that have diverged between zebrafish and *Drosophila* to unravel how dysregulation in this signaling pathway results in developmental abnormalities^19,20^ and diseases such as cancer ^21^. Homology between zebrafish and *Drosophila* is evident through the major regulators and their functions. In zebrafish, BMP ligands, BMP2/7, are homologous to *Drosophila* Dpp/Scw, while Chd, a homolog of *Drosophila* Sog, acts as an inhibitor of the signal. Even with this level of conservation, when Chd was introduced to *Drosophila*, Chd failed to fully rescue *Drosophila* sog^−/−^ embryos to levels of WT BMP ^22^, suggesting an evolutionary divergence in the mechanism. *Drosophila* Sog has been thought to play a paradoxical role as both an inhibitor and a promoter of BMP^23–25^ signaling, while Chd, in zebrafish, does not exhibit this dual function. In addition to this biochemical difference between *Drosophila* and zebrafish, there are also substantial differences in the spatial and temporal scales. *Drosophila* has a smaller embryo, less than half the size of the zebrafish embryo. In terms of time scale, the *Drosophila* Dpp gradient is established to pattern the DV axis in approximately 1 hour after cellularization ^16,26^ while zebrafish BMP gradient develops gradually in about 2-3 hours^17,27^. The establishment of the BMP gradient depends on the balance of all these factors: the biochemical reactions, embryo geometry, and the length and time scales of the systems.

Multi-objective optimization (MOO) is an approach that allows model parameter optimization using multiple datasets by defining the different datasets as different objectives^28,29^ and allows researchers to consider multiple conflicting objectives simultaneously. Different datasets may vary in accuracy and importance in different scales, which are often unknown; therefore, they cannot simply be aggregated into a single objective without proper weights. Unlike single objective optimization, MOO provides a set of compromise solutions, known as the Pareto front, that reveals all trade-offs between objectives. Although MOO provides a desirable approach for integrating all experimental data without prior knowledge of precision, its high computational cost has limited its use in biological systems. Areas that have applied MOO in biology are limited to parameter estimation ^30–36^, optimal control ^37–40,^ and experimental design ^41–43^. Most of these approaches use simulated data instead of real experimental data to demonstrate the methodology. In this work, we use a mathematical model analysis and multi-objective optimization strategy to study and understand the minimum requirements on the biophysical parameters that balance the known geometry, length, and time scale differences between *Drosophila* and zebrafish to produce observed BMP gradient data. We investigated how a single model could fit different systems by varying the model parameters to determine the homology (conservation) and also identify the diversification of the BMP pathway in both systems. Mathematical models of BMP signaling have provided some valuable insights into the gradient formation of BMP and how it might be regulated by regulators^4,44–49^. These models describe how the morphogen concentrations change over time and space through diffusion and chemical reactions such as synthesis, degradation, association, and dissociation with other factors (such as antagonists). We use a simple core model that captures the classical interactions of the homologous BMP regulators in both *Drosophila* and zebrafish. This model relies on the balance between diffusion, reaction, and degradation of a locally produced signal^1,45,50^. The signal can be realized locally by protein synthesis and could be degraded through proteolysis or receptor-mediated endocytosis, while diffusion is a result of ligand internalization and recycling. The interaction between these processes generates different variations of the model, such as free diffusion (diffusion-decay) models^45,50–53^, hindered diffusion (reaction-diffusion) models^45,54–58^, and facilitated diffusion (shuttling) models^4,24,44,46,47,57,59,60^. It has been shown that other regulatory processes are essential to produce the observed BMP signal in both *Drosophila* and zebrafish. For *Drosophila*, positive feedback in response to pMad signaling is needed to further concentrate the surface localized Dpp/Scw at the dorsal midline^26^. In zebrafish, other inhibitors, Noggin (Nog) and Follistatin, in addition to Chd, are essential for the establishment of some ventral cell fates in the early embryo^61^. Our work aims to investigate whether a single mechanism that is homologous exists between *Drosophila* and zebrafish to produce observed phenotypes.

Pareto fronts provide valuable insights into the relationships between objectives in a system, models or datasets. The shape of the Pareto front reveals how objectives interact: a convex shape illustrates a lot of indistinctness in the objectives, with a steeper shape indicating the potential for a single compromise solution that adequately satisfies both objectives^62^. In contrast, a linear Pareto front suggests that the model is not capable of meeting both objectives simultaneously, revealing a degree of both indistinctiveness and distinctiveness of the objectives. A concave shape, on the other hand, suggests that the objectives are completely distinct and cannot be optimized simultaneously. Pareto fronts can also be compared for dominance using the hypervolume quality indicator (also known as S-metric or Lebesgue measure)^63–65^. While these tools are available to compare Pareto fronts, they have not been used in biology because of their computational expense. In addition, little work has been done to address uncertainty associated with frontiers in multi-objective optimization, as most uncertainty propagation methods have been developed for single-objective problems. Extending these methods to multi-objective optimization remains computationally expensive and unexplored. In this work, we introduce a novel concept of propagating uncertainty to a Pareto front to account for data uncertainty. Additionally, we extend the concept of Pareto optimality by introducing the Utopian front, a new frontier that relaxes constraints to identify the minimum set of design variables required to be different to achieve a solution that converges to the utopian point that occurs with no constraints and complete independence of design variables in model optimization. This extension of Pareto optimality provides a powerful tool to analyze and identify limitations in complex systems with competing objectives.

In a recent study, we investigated how the BMP/Smad pathway balances multiple systems-level behaviors—such as response speed, noise filtering, and sensitivity—across various cellular contexts with the Pareto front analysis ^66^. Despite the pathway’s core connectivity remaining largely unchanged, our findings showed that non-conserved parameters (NCPs), including protein concentrations in the Smad pathway and especially nuclear phosphatase, can be tuned to optimize different performance objectives. However, the Smad pathway alone cannot simultaneously maximize all objectives. In this study, we extend this multi-objective optimization perspective to an organismal scale—contrasting Drosophila and zebrafish—and demonstrate how key extracellular parameters also shape BMP signaling outcomes across species.

## Methods

### Data collection and calibration

We aggregated measurements for BMP signaling from literature sources and previously published work in various formats for both Drosophila and zebrafish. The primary data of interest included the intensities of the BMP signal transducer, pMad in *Drosophila*, and pSmad in zebrafish, measured across the D-V axis of their respective embryos. A summary of the data is provided in Table 1. The available data were (1) quantitative (pMad/pSmad intensities), (2) semi-quantitative (images that reflect the spatial distribution of the signaling molecules), and (3) qualitative (descriptive interpretations of the observed phenotype of wild-type and mutant embryos). To ensure consistency, all collected data were calibrated against internally processed experimental data and scaled based on intensity and geometry for both systems^67–69^ for model fitting (*All data available in the supplement). Drosophila* embryo data were collected from dorsal-orientated embryo images. A rectangular region, measuring 10% the width of the embryo was then cut in the middle of the A-P axis and averaged to get a one-dimensional pMad distribution along the DV axis. Since the dorsal view image only shows half the embryo, the BMP gradient is positioned from the dorsal to the ventral lateral regions of the embryo. The position of the high BMP signal is labeled position x = 0, dorsal midline for the *Drosophila* embryo, and the ventral midline for zebrafish.

**Table 1.**
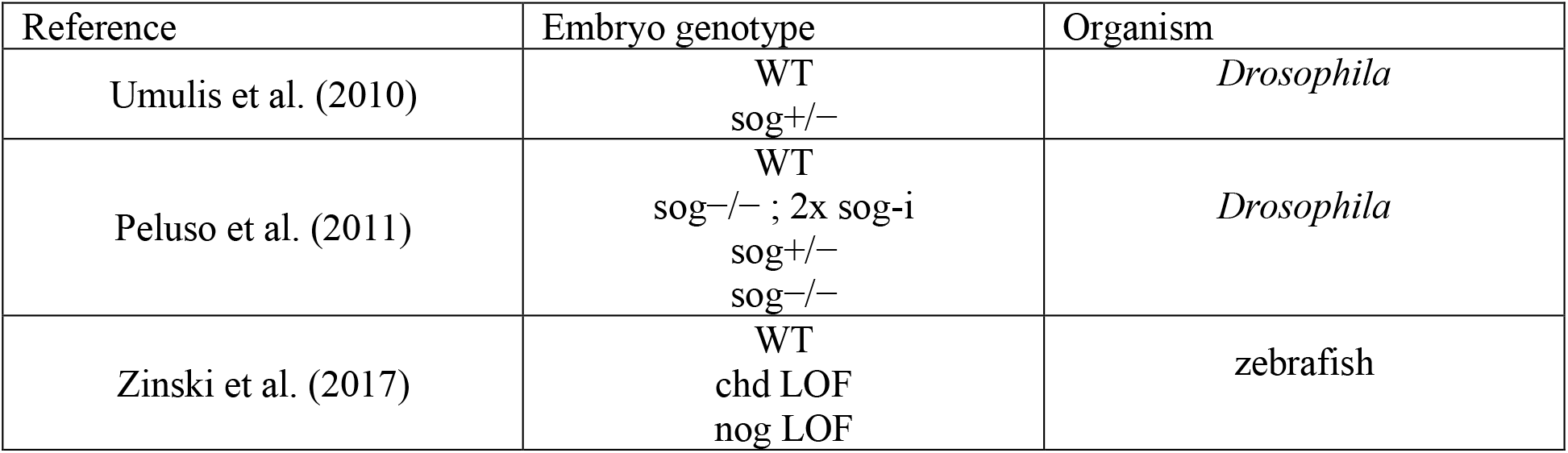
Data extracted from published literature.

We built a single core model to determine parameters wherein one model could fit both systems and if not, identify the minimum number of changes in the parameters needed to fully fit both systems. We used the simplest core model shared between *Drosophila* and zebrafish to generate a BMP profile to fit against the experimental data. The model considered here is a simplified version of our previously published BMP models ^70^. This mechanism relies on the diffusion and degradation of a locally produced signal^1,50,71,72^. To form an appropriate simplified model for both *Drosophila* and zebrafish systems, we applied a set of assumptions: (i) reactions follow first and second-order kinetics, (ii) the system is represented in one dimension, (iii) BMP heterodimer formation is rapid and negligible, allowing us to model only a single molecule. For the analysis, we focused on key variables that control BMP inhibitor expression strength and the boundaries of the Chordin/Sog expression and in situ expression data.

Based on these assumptions, we formulated a system of ordinary differential equations (ODE) to describe the interactions of regulatory factors. Equations 1, 2, and 3 represent a one-dimensional line of cells oriented along the dorsal-ventral axis where long-range diffusion is approximated using the finite difference method, shown in Equation 4. We applied no-flux boundary conditions to both of the systems, and all factors were initially set to a null concentration (zero value). The complete set of equations for the simple model is as follows:

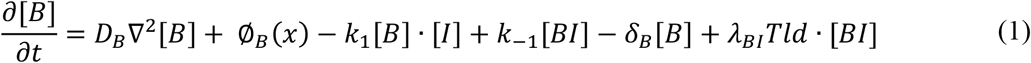

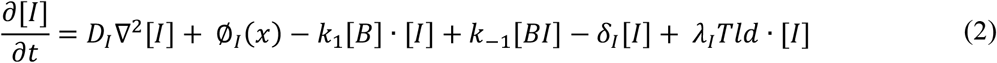

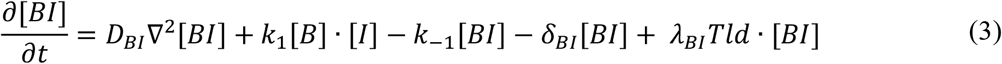

where [B], [I], and [BI] represent the concentrations of BMP, the inhibitor, and the BMP-inhibitor compound, respectively. The model is parameterized by D_j_, the diffusion rate, δ_j_, the decay rate, and φ_j_ (x), the production rate of protein j where j = {[B], [I], [BI]}. BMP binds to the inhibitor with a rate of k_1_ and unbinds with a rate of k_−1_, Tld cleaves BMP bound to the inhibitor with a rate of λ_BI_ and cleaves free inhibitor with a rate of λ_I_. The inhibitor, [I], is Sog in Drosophila and Chd in zebrafish.

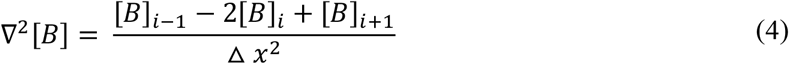

The solution depends on the balance of time scales, diffusion, chemical reactions, and embryo geometry. We assume that the fundamental diffusion and chemical reaction equations for both systems are similar, as described in Equations 1, 2, and 3. However, since the shape of the BMP signal differs greatly between *Drosophila* and zebrafish, we code these differences with variations in the biophysical parameters, as well as distinct geometries, length scales, and timescales of the two systems. These known differences are summarized in Table 2.

**Table 2.**
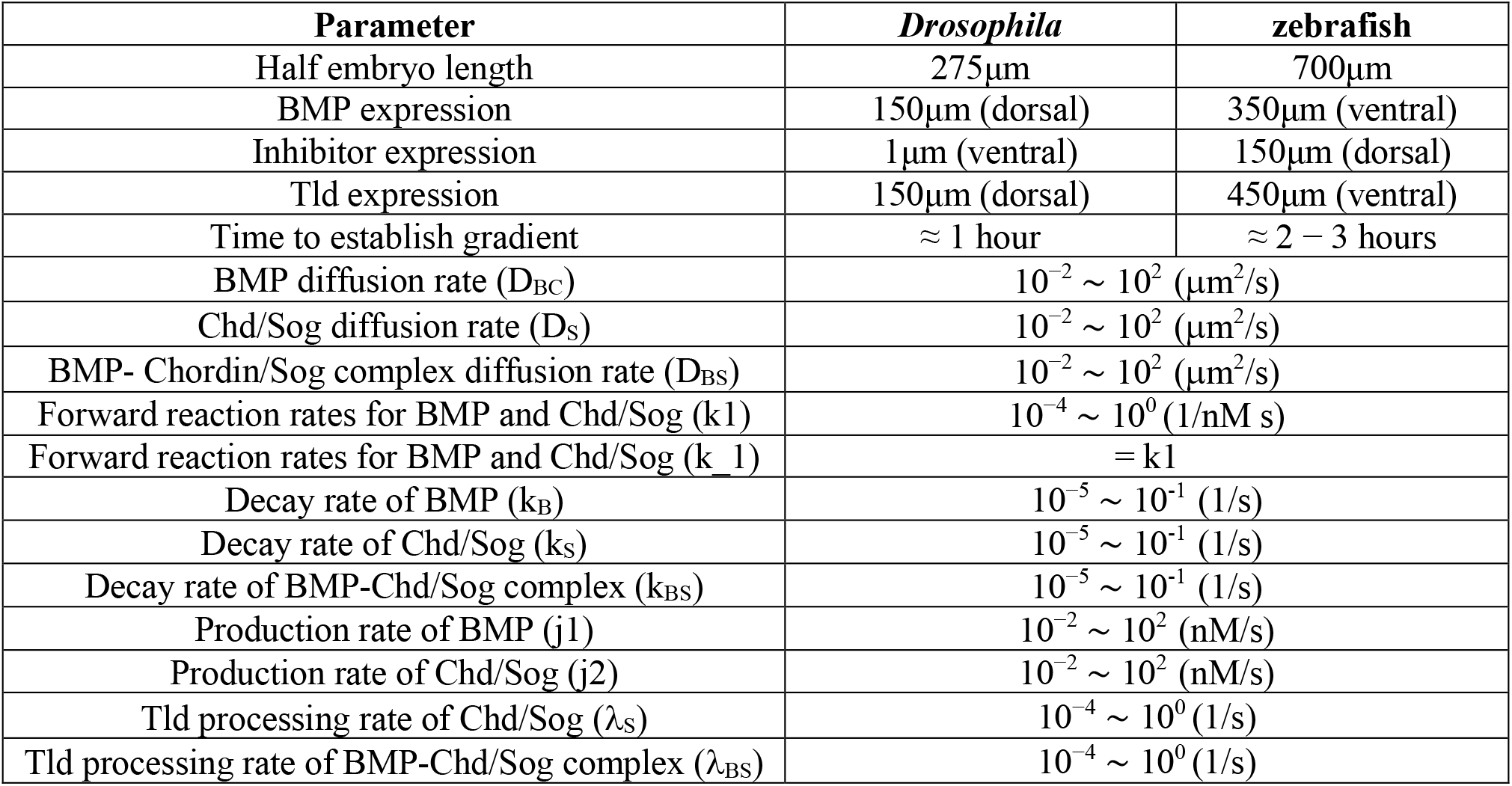
Parameters that specify the length and time scales of Drosophila and zebrafish.

To evaluate the system across a broad parameter space, we conducted an initial search through the computational model-based screen of over 200,000 combinations of biophysical parameters of the major extracellular BMP modulators. We applied the Latin Hypercube Sampling (LHS) strategy wherein 11 independent parameters were selected from a uniform distribution in log space that spanned four orders of magnitude within the physiological range for each parameter. The parameter ranges used for sampling are listed in Table 2. For each sampled parameter set, we first performed an initial wild-type simulation; the model was then re-simulated with Chordin production set to zero to simulate the BMP signaling gradient in a *sog*(−/−) *Drosophila* mutant, and Chordin loss-of-function (LOF) scenario in zebrafish cases. All computations were performed using Purdue’s Supercomputing Clusters.

### Parameter sensitivities

A Latin Hypercube Sampling Partial Rank Correlation Coefficient (LHS/PRCC) sensitivity analysis is employed to analyze how the objective function varies with model parameters to identify the most relevant parameters that influence data fitting. LHS/PRCC is particularly useful as it explores the entire parameter space of a model with a minimum number of model simulations ^73^. PRCC is based on linear regression analysis and evaluates correlation coefficients between model inputs and outputs. The correlation coefficient is normalized between −1 and 1, which allows comparisons between model inputs. Herein, we focused specifically on the sampled points that are close to the Pareto front (Pareto local) to reduce the bias to the sensitivity caused by significant changes in the objective due to ill-fitting parameters. The sensitivity of the objective is analyzed on the points that are within the 95% confidence interval of the Pareto front when the uncertainty is propagated from experimental data, as described in detail in Box 1.

#### Box 1

**Pareto Front uncertainty**

Uncertainty in measurements propagates through objectives optimization, affecting Pareto fronts. To quantify this uncertainty in Pareto fronts, we developed a method based on Normalized Root Mean Square Error (NRMSE) as the objective function. The experimental data were defined as 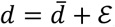, where 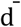 is a vector of mean data points and *ε* is a vector of normally distributed random variable, *ε*~*𝒩*(*μ, V*). The objective function is defined as:

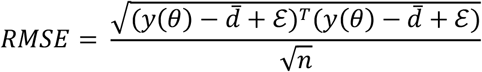

where *y*(*θ*)is the model simulated by parameter vector, *θ* and *n* is the number of data points in a dataset. Using a Taylor series, we linearized the objective function around the mean of the data, 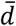, where *ε* = 0:

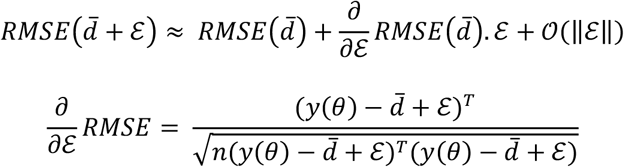

Therefore, the linearize objective around the mean data is,

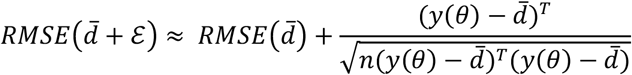

which could be simplified to,

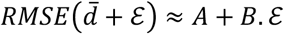

where *A, B* are a constant scalar and vector, respectively that are independent on *ε*. Since *ε* is normally distributed with mean, *μ* = 0 and covariance matrix, *V*, we approximated the distribution of 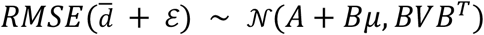.

We introduced a novel concept of the Utopian frontier to compare the compromise front between Drosophila and zebrafish system, which is described in Box 2.

#### Box 2

**Utopian Front**

We introduce a novel concept of Utopian frontier which extends the concept of Pareto frontiers in multi-objective optimization. The Utopian frontier is defined as a set of utopian solutions created by selectively optimizing a subset of the unknown parameters while holding the remaining parameters fixed at their Pareto optimal values. Therefore, the Utopian frontier is derived from an existing Pareto frontier. The set of Pareto points is attained by solving a multi-objective optimization problem expressed as:

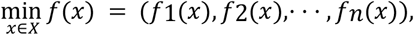

where *fi* ∈ *R*^*n*^ are multi-objectives functions of x constrained within a design space *X*; *G*(*x*) ≤ 0 and *X* ∈ *R*^*d*^. A Pareto front consists of points, *x*^*^ ∈ *X* that satisfy:

1. *fi*(*x*^*^) ≤ *fi*(*x*) for all indices *i* ∈ 1, 2, …, *n* (strict dominance) and
2. *fj*(*x*^*^) < *fj* (*x*) for at least one index *j* ∈ 1, 2, …, *n* (weak dominance).

The set of all Pareto optimal design variables is denoted as *X*^*^. A global utopia point is an idealized, theoretical infeasible point that simultaneously optimize all objectives.

As demonstrated in Figure 2, for each *x*^*^ ∈ *X*^*^, we re-optimized the objectives by varying only a subset, *x*_*S*_ ∈ *x*^*^, while holding the remaining complement subset *x*_*S*_′ constant. reduces the optimization search space to, *S* = {*x*_*S*_} ∈ *R*^*q*<*d*^ while the complement subset, *S*′ = *{x*_*S*_′} ∈ *R*^*d*−*q*^ is treated as equality constraints. The new multi-objective optimization problem is then defined over the subset, *S* for each Pareto solution, *x*^*^ = (*x*_*S*_, *x*_*S′*_) as follows:

**Figure 2:**
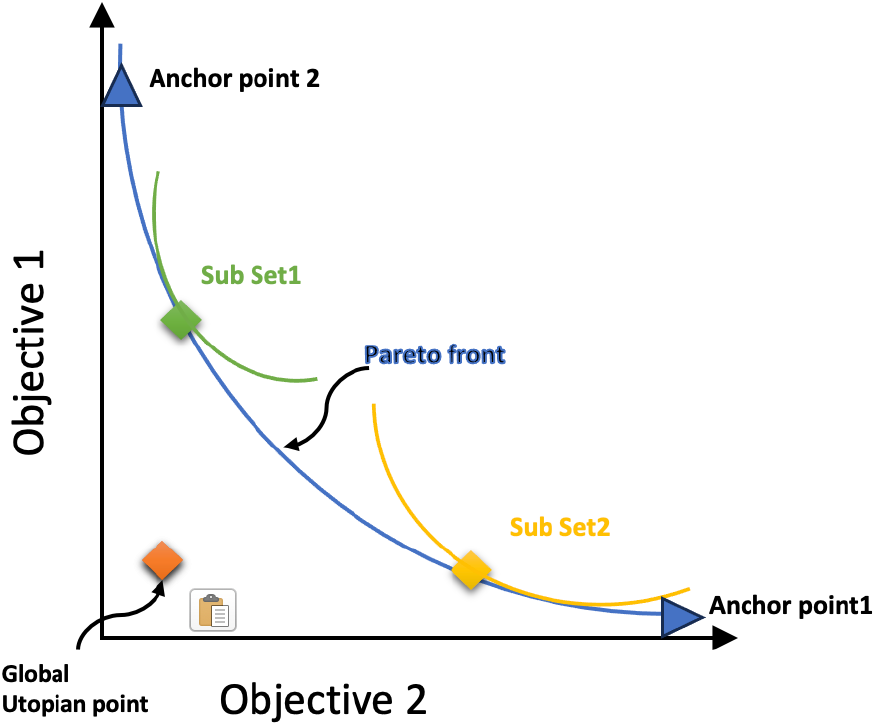
An illustration of the Utopian front. The Pareto front (blue curve), Utopian fronts (green and yellow curves) examples, and the Global Utopia Point (orange diamond) are visually shown. The Pareto front captures the optimal trade-off between two objectives, while Utopian fronts are generated by optimizing different subsets of parameters. Subset 1 (green curve) and Subset 2 (yellow curve) demonstrate distinct optimization subsets anchored at different Pareto solutions.

**Figure 3.**
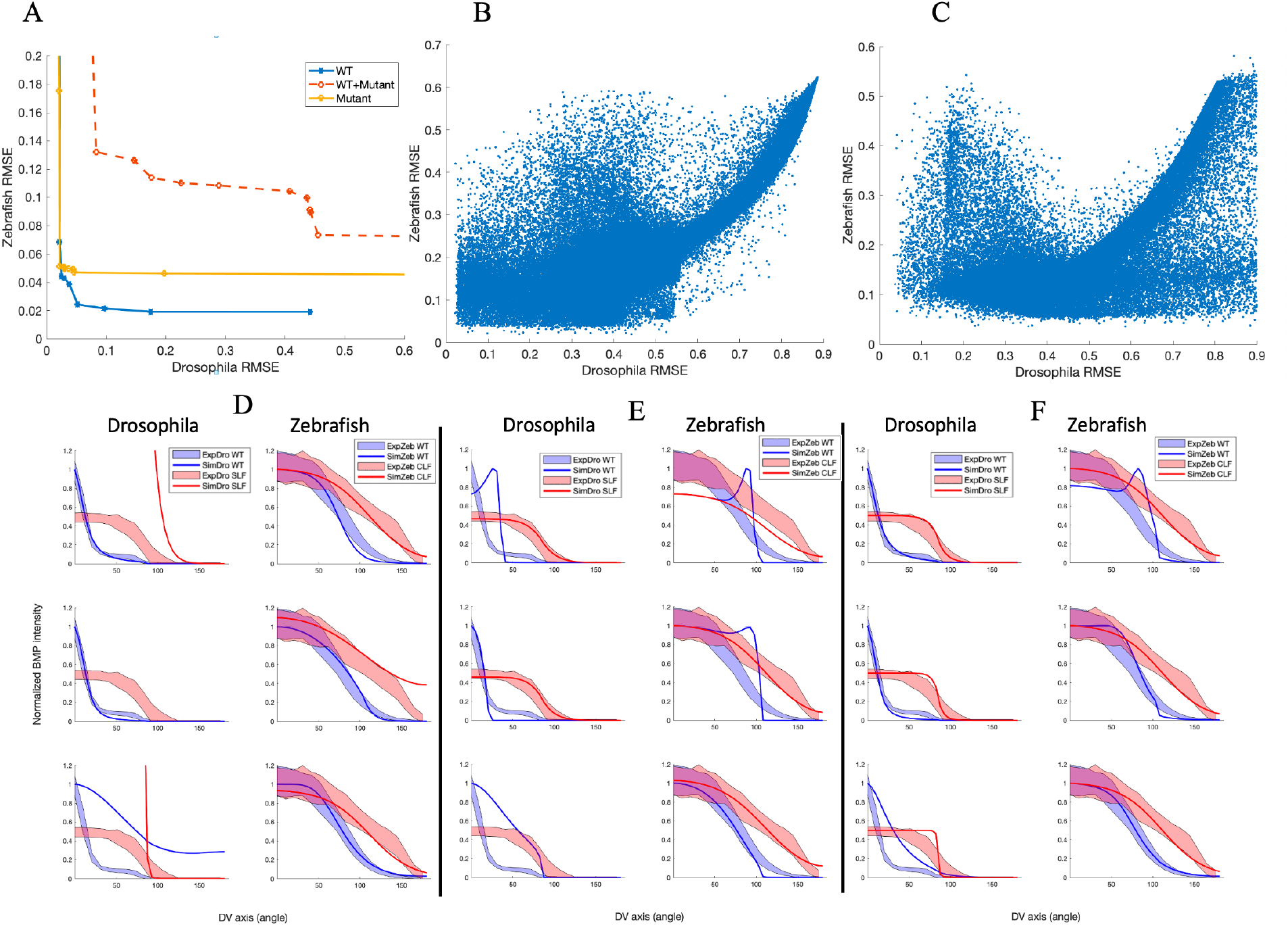
Core model Pareto Front analysis. (A) Pareto front generated by fitting the core model to WT, mutant(sog(−/−)/Chd LOF), and both WT and mutant data. (B) Core model simulation results space for WT fitting (C) Core model simulation results space for sog(−/−)/Chd LOF fitting. Corresponding model simulations with Drosophila anchor point (top panel), best compromise point (middle panel), and zebrafish anchor point (lower panel) are compared to (D)WT experimental data, (E) sog(−/−)/Chd LOF data, And (F) both WT and mutant data. ExpDro and ExproZeb indicate experimental data for Drosophila and zebrafish, respectively, while SimDro and SimZeb indicate simulations for Drosophila and zebrafish, respectively. Mutant data indicate sog(−/−) mutant in drosophila data and Chd LOF in zebrafish case for simplifying.

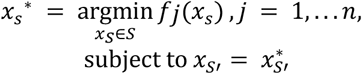

where *j* are the n objective functions and 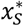 is the set of solutions creating a new Pareto frontier that is tangent to the original Pareto front at the point, *x*^*^. The Utopian point of the new Pareto frontier is the ideal point which could simultaneously optimize all objective functions, defined by *x*_*u*_ as follows:

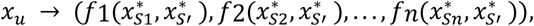

where 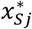 are the anchor points for each *j*^*th*^ objective. The collection of all non-dominated Utopia points, *x*_*u*_ ∈ *X*_*u*_ (Utopian front) solved along the original Pareto frontier, *x* ∈ *X*. The Utopian front lie between the original Pareto front and the global utopia point. The convexity and proximity of a Utopian front from the Pareto front reveal the relationship between parameter subsets and the trade-offs among objectives.

- A static Utopian front, invariant to the Pareto front, suggests that optimizing different parameter subsets does not improve any of the objectives.
- A Utopian front close to the global utopia point suggests that varying certain optimal values of the subset variables of each objective can improve all objective functions.

## Results

### Fixed parameter common core model fails to simultaneously fit Drosophila and zebrafish systems under the same biophysical conditions

To evaluate whether the simplest core model, described in Equations 1, 2, and 3, could adequately describe both *Drosophila* and zebrafish BMP regulatory systems, we generated Pareto solutions by fitting the model to both systems with WT and mutant (*sog*(−/−) in *Drosophila* and Chd LOF in zebrafish) data simultaneously. The Pareto solutions obtained from fitting the WT data along and *sog*(−/−)/Chd LOF only and fitting the WT with mutant are shown in Figure 3A, sampled from the parameter space in Figure 3B and 3C. The Pareto front generated from WT data only exhibits high convexity, suggesting that there is a common set of biophysical parameters that fit WT *Drosophila* and zebrafish experimental data. The best compromise solution demonstrates high fitness to the WT data (blue line in Figure 3A) with objective function penalties closer to zero (0.0242 and 0.0524 for *Drosophila* and zebrafish systems, respectively). However, this result seems to suggest a possible simple conserved mechanism between the two systems. Similarly, when fitting only mutant data (*sog*(−/−)/Chd LOF), the best-compromise solution for model simulations aligned well with experimental observations (yellow line in Figure 3A), yielding residuals of 0.0367 and 0.0498 for *Drosophila* and zebrafish systems, respectively.

However, while the model performed well when fitting WT (Figure 3D) or mutant data (Figure 3E) separately, it failed to reproduce both WT and mutant datasets simultaneously (Figure 3F). When fitting both WT and mutant experimental data simultaneously, the resulting Pareto front displayed less convexity compared to the WT-only or mutant-only Pareto front. This suggests an underlying divergence in BMP signaling mechanism between *Drosophila* and zebrafish that is not captured by the core model. Although the model’s ability to fit mutant data improved, the best compromise solution still failed to fully capture the BMP signaling mechanism in *Drosophila*. Additionally, the model was insufficient in discriminating between WT and mutant data, indicating that additional regulatory mechanisms or parameter variations may be required to accurately model both species under the same biophysical conditions.

### Chordinlike variants shift *Drosophila* system closer to zebrafish while preserving shuttling

In a key rescue experiment, ‘Chordinlike’ variants introduced into *sog(−/−)* Drosophila embryos significantly broadened the steep BMP gradient observed with endogenous Sog ^22^. Notably, Peluso et al.^22^ found that while Sog requires BMP for efficient cleavage by Tld, Chd can be cleaved by Tld in the absence of BMP. This fundamental difference in cleavage specificity helps explain how the Drosophila gradient can be shifted to resemble that of vertebrate embryos while maintaining a degree of BMP shuttling ^74–76^. To implement this phenomenon, we specified the Tld cleavage rate of ‘Chordinlike’ Sog was specified to be slightly higher than that of Indigenous Sog and investigated whether the broader BMP gradient observed with ‘Chordinlike’ Sog could indicate a shared regulatory mechanism between *Drosophila* and zebrafish. The resulting Pareto front generated by fitting the model to “Chordin-like” data dominates the Pareto front of the indigenous system (Figure 4A). The differences in the Pareto fronts are particularly pronounced along the trade-off regions, suggesting that with the “Chordin-like” Sog, the *Drosophila* system shifts towards a BMP mechanism similar to zebrafish. To ensure that the observed differences in Pareto fronts were not simply due to propagated uncertainty, we applied an uncertainty propagation technique. Figures 4A, 4C, and 4E illustrate how the uncertainty propagates from experimental measurements to the objective space for both models. The error bars represent 95% confidence intervals, and their non-overlapping nature confirms that the Pareto frontier differences are not simply due to experimental. This result, therefore, confirms that the observed shift in BMP regulation is due to the biophysical difference between endogenous Sog and “Chordin-like” Sog variants.

**Figure 4.**
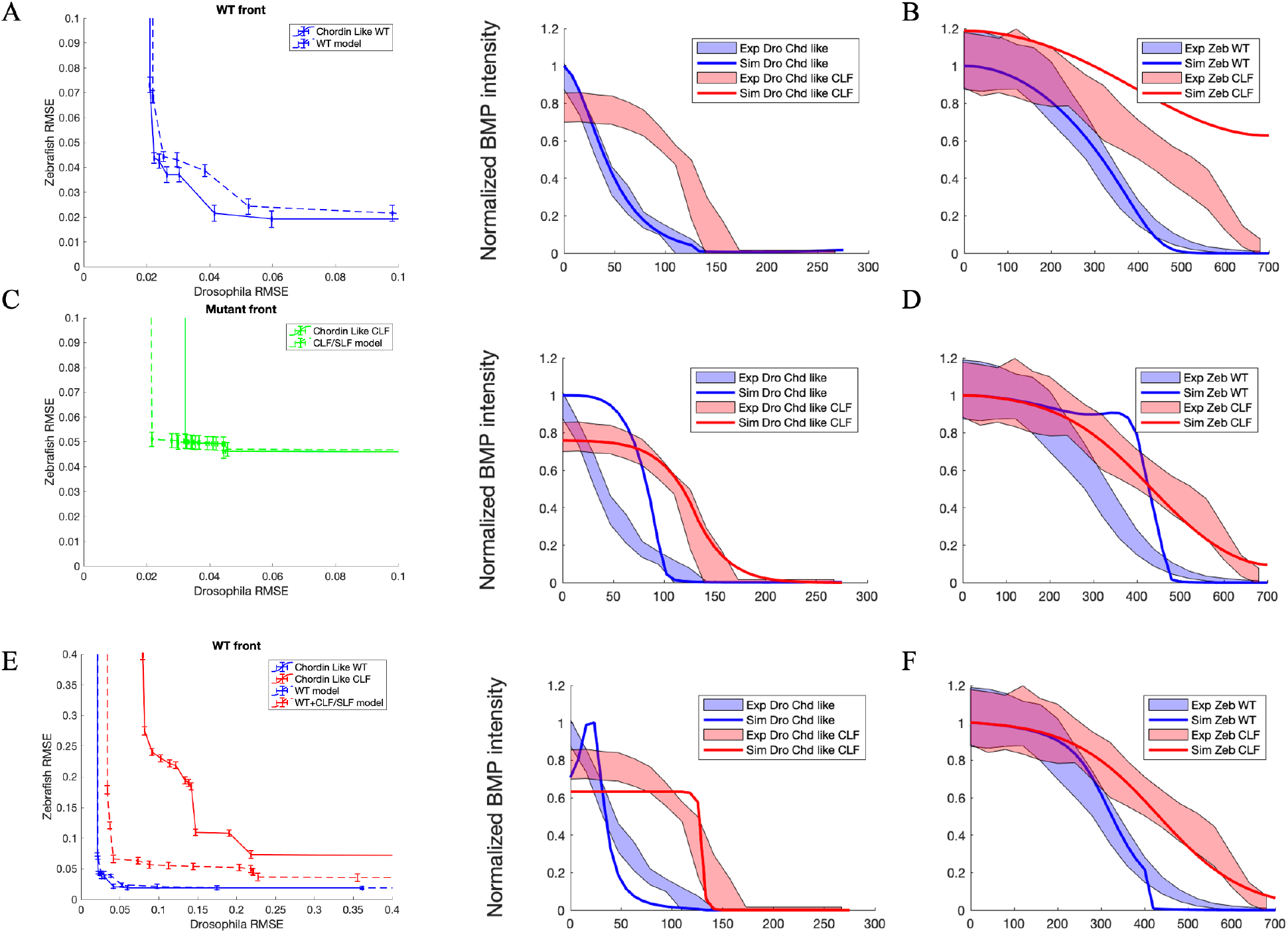
Comparison of indigenous and ‘Chordin-like’ Core Model Pareto fronts. The ‘Chordin-like’ Pareto front dominates the indigenous front, demonstrating that ‘Chordin-like’ data shifts the Drosophila BMP mechanism to resemble that of zebrafish for both the WT fitting (A), ‘Chordin-like’ sog (−/−)/ Chd LOF mutant fitting (C) and WT+ ‘Chordin-like’ sog (−/−)/ Chd LOF mutant fitting (E). Corresponding model simulations of the best compromise point for both Drosophila and zebrafish are compared to ‘Chordin-like’ experimental data. (B) Compromise solution for ‘Chordin-like’ Drosophila data vs WT zebrafish data. (D) Compromise solution for ‘Chordin-like’ Sog(−/−) mutant Drosophila data vs Chd LOF zebrafish data. (F) Compromise solution for ‘Chordin-like’ +’Chordin-like’ Sog(−/−) mutant Drosophila data vs WT+Chd LOF zebrafish data.

Surprisingly, despite the biophysical differences between Sog and Chd revealed by the “Chordin-like” experiment, BMP shuttling in *Drosophila* remained intact. As shown in Figure 4D, BMP movement toward the dorsal midline was still observed, indicated by a reduced BMP gradient in *sog(−/−)* embryos compared to WT. Although shuttling was preserved, its strength was diminished. The WT BMP gradient remained broader, and the *sog(−/−)* simulation shows only about a 20% decrease. Furthermore, the slight increase in the Tld processing rate of free Sog corroborates previous findings that Sog processing requires BMP, whereas Chd can be cleaved by Tld independently of BMP^22^. Finally, the Pareto front generated by the “Chordin-like” mutant (Figure 4C) aligns more closely with Drosophila *sog(−/−)* data than with zebrafish Chd LOF data, underscoring the unique biochemical properties that differentiate Sog from Chd.

### Chordin/Sog-related parameters shape the Utopian front between Drosophila and zebrafish

To determine the influence of individual parameters on the Pareto front, we evaluated how the 11 independent biophysical parameters affect the solution space and Utopian fronts. We examined 200,000 simulation results in RMSE space for both *Drosophila* and zebrafish under both WT (Figure 5B) and mutant conditions (Figure 5B), with a color scale reflecting the specific range for the individual parameter. We observe that specific parameters, particularly those linked to Chordin/Sog activity, play a prominent role in shaping the Utopian front. In the WT scenario (Figure 5A), the diffusion rates of BMP (D_B_), Chd/Sog (D_S_), and the BMP–Chd/Sog complex (D_BS_), as well as the production rates of BMP (J_1_) and Chd/Sog (J_2_), tend to cluster in regions closer to the Utopian point. A similar trend is observed under sog(−/−)/Chd LOF conditions but with an additional emphasis on λ_BS_ (the Tld processing rate of the BMP–Chd/Sog complex), which becomes more pronounced in mutant simulations. We further evaluated how changes in these parameter ranges shift the Utopian front (Figure 6). Notably, increasing the diffusion rate of Chd/Sog (D_S_) consistently draws the Utopian front towards an optimal utopian point (Figure 6A). This effect is more pronounced when D_S_ falls within the range of 10^−2^ to 10^−1^ (μm^2^/s).

**Figure 5.**
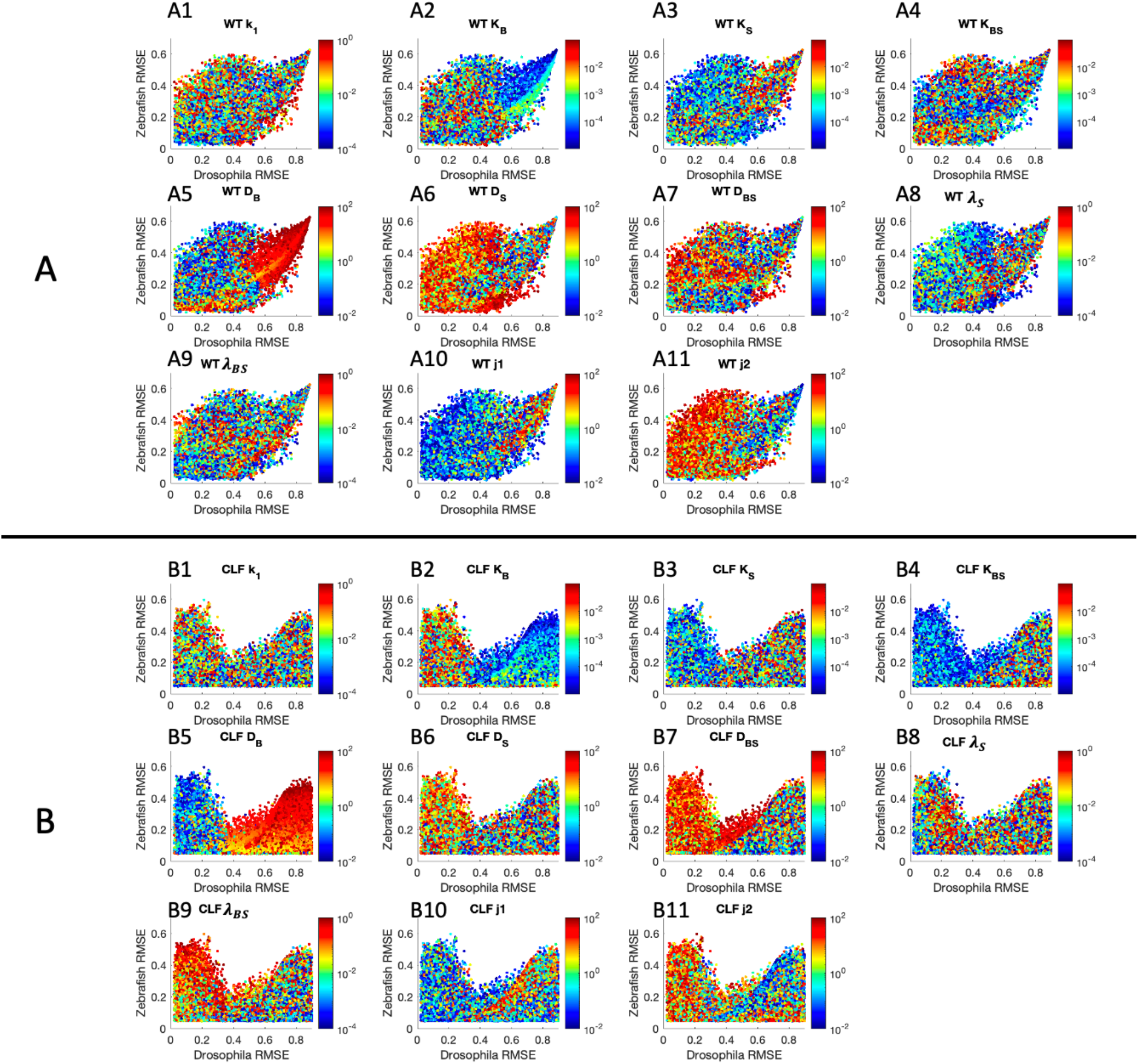
Parameter trend analysis for all the simulation results against WT and Chd/Sog(−/−) fitting. Figure group A shows the parameter tendency in Utopian space against WT fitting for 11 independent parameters, including A1, k_1_(forward reaction rates for BMP and Chd/Sog); A2, K_B_ (Decay rate for BMP); A3, K_S_ (Decay rate for Chd/Sog); A4, K_BS_ (Decay rate for BMP-Chd/Sog complex); A5, D_B_ (Diffusion rate for BMP); A6, D_S_ (Diffusion rate for Chd/Sog); A7, D_BS_ (Diffusion rate for BMP-Chd/Sog complex); A8, λ_S_ (Tld processing rate of Chd/Sog); A9, λ_BS_ (Tld processing rate of BMP-Chd/Sog complex); A10, J1(Production rate of BMP); A11, J2 (Production rate of Chd/Sog). Group B shows the parameter tendency in Utopian space against Chd/Sog(−/−) fitting for 11 independent parameters, respectively. Notably, the RMES range is limited to [0,0.6] for Zebrafish fitting and [0,0.8] for Drosophila fitting for better visualization.

**Figure 6.**
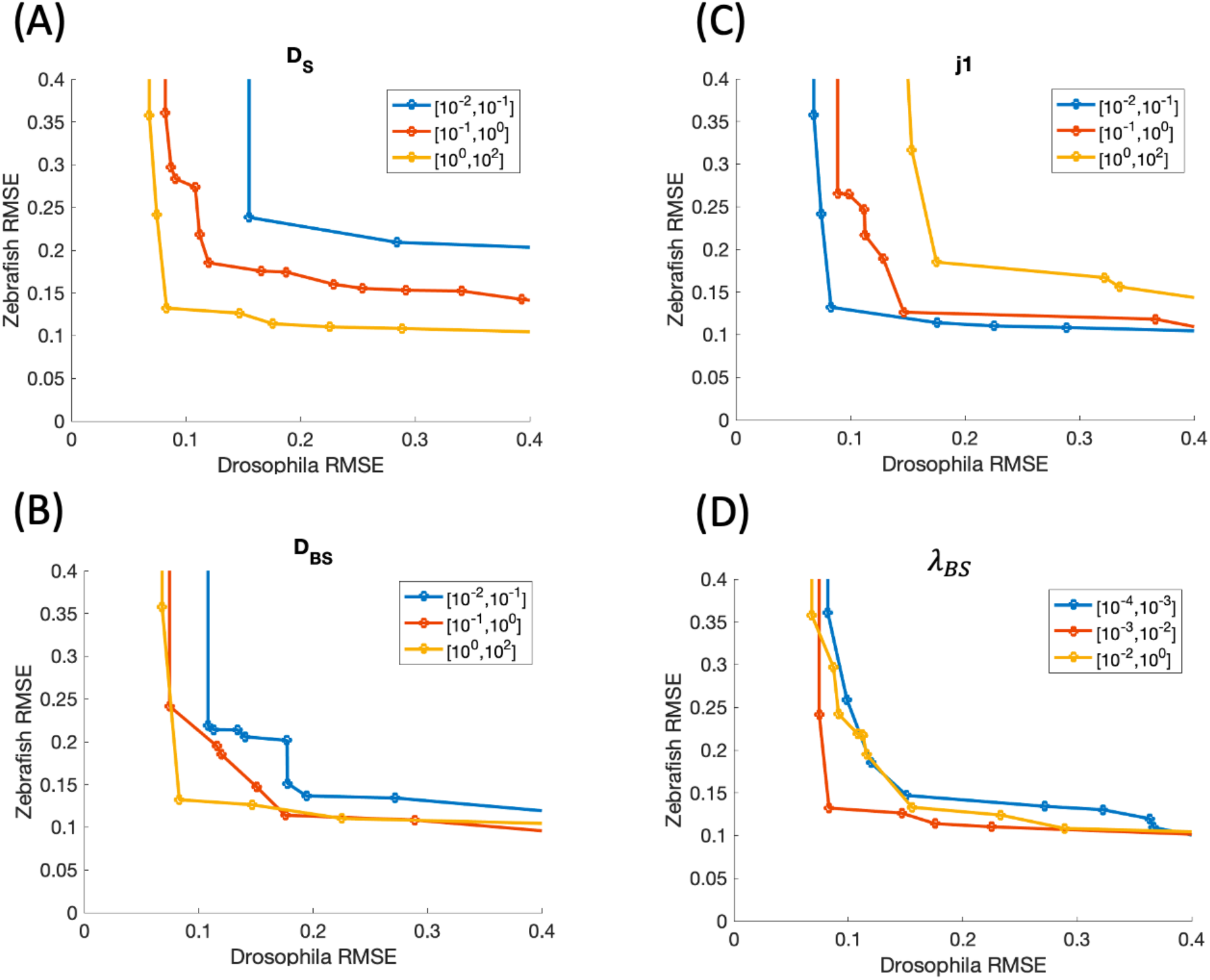
Utopian front of WT + Sog/Chd(−/−) fit for the individual parameters in specific ranges. D_S_ (Diffusion rate for Chd/Sog); A7, D_BS_ (Diffusion rate for BMP-Chd/Sog complex); J1(Production rate of BMP); λ_BS_ (Tld processing rate of BMP-Chd/Sog complex)

Additional simulations using the “Chordin-like” variant (Supplementary Figures S1) show analogous parameter trends and confirm that elevated diffusion rates and processing factors shift the Utopian front toward an optimal region, reinforcing our primary findings. These findings suggest that while certain parameters share broad optimal ranges across the *Drosophila* and zebrafish, others—especially those related to Chordin/Sog processing and diffusion—may exhibit species-specific constraints, suggesting the evolutionary divergence in BMP regulatory mechanisms between the two species.

A notable finding of our analysis is the identification of specific biophysical parameters that act as critical “switches” governing BMP gradient formation in Drosophila and zebrafish. These parameters seem to diverge between species, driving system-specific solutions along the Utopian front. Among these key parameters, the diffusion rate of Chordin/Sog (DS) (Figure 5 and 6A), the diffusion rate of the BMP– Chd/Sog complex (D_BS_) (Figure 5 and 6B), and the Tld processing rate of the BMP–Chd/Sog complex (λ_BS_) (Figure 5 and 6D) stand out as important. These parameters exhibit distinct optimal ranges between Drosophila and zebrafish, suggesting that evolutionary shifts in their values contributed to divergent dorsal-ventral patterning strategies in these two species. Further Utopian front analyses for other biophysical parameters are shown in Supplementary Figure S2. The differences in these parameters provide a more apparent mechanistic explanation for why substituting Chordin in Drosophila fails to fully rescue sog mutant phenotypes. More generally, these results highlight how subtle biophysical alterations in morphogen diffusion and processing can drive significant variations in tissue patterning across species. Interestingly, the parameters that most significantly shift the Pareto front align with those identified as key sensitivity factors in our previous 2D BMP model study. Based on these findings, we plan to extend our research to explore species-specific parameters within this system.

### Relaxed model results show Utopian fronts alluded to a possible divergence in BMP mechanism between *Drosophila* and zebrafish

The BMP gradient relies on the balance of geometry, biophysical interactions, and conditions of the systems to establish observed BMP gradients in both *Drosophila* and zebrafish. Herein, we take a step-by-step investigation to determine the minimum combination of biophysical interactions and parameter conditions needed to generate the differences in the mechanisms. A Utopian front outperforms the original Pareto front, suggesting that allowing the biophysical parameters to differ between *Drosophila* and zebrafish improved the model’s ability to fit both sets of data. To identify the influential parameters combination that improves fit for both systems, we tested our model by introducing a relaxation scheme based on our prior sampling of model parameters. This approach introduces additional degrees of freedom (DoF) in the systems by allowing subsets of shared parameters to vary between *Drosophila* and zebrafish. Specifically, in the one-degree scenario, a single parameter was allowed to vary between the Drosophila and zebrafish models, while all others remained constant and identical in both systems. This process was repeated for all 11 parameters. The two-degree scenario involved allowing two parameters to vary between the systems, producing 55 unique combinations, while the three-degree scenario produced 156 possible combinations of relaxed parameters. These relaxation tests were conducted on the top of the initial screen of 200,000 parameter sets, where both systems shared identical parameters (considered a relaxation degree of zero). Our results revealed a more prominent Utopian front as the DoF increased in WT only (Figure 7A), mutant only (Figure 7B), and WT + mutant conditions (Figure 7C). However, while the *Drosophila* mutant Utopian front moved towards the Utopian point with increased relaxation, the zebrafish mutant fitting showed little improvement, implying that the model has a limited capacity to capture mutant conditions in the zebrafish system (Figure 7B).

**Figure 7.**
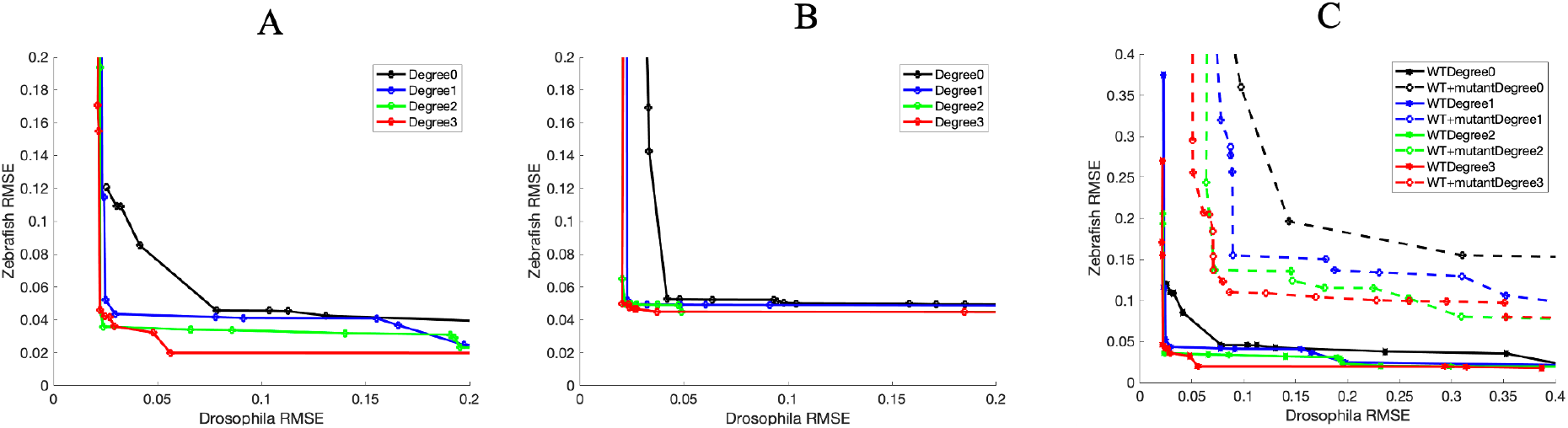
Pareto Front for released model against (A) WT fit, (B) Sog (−/−)/Chd LOF fit, (C)Comparison between WT fit and WT + Sog (−/−)/Chd LOF fit fronts. Degree 0, based on 10,000 simulations in Monte Carlo sampling in whole parameter space. Degree 1 releases one parameter (total of 11 parameters) in the parameter set of degree 0, which can vary between the Drosophila and Zebrafish models. Degree 2 releases two parameters, and Degree 3 releases three parameters.

## Conclusion

Our work highlights the use of multi-objective optimization (MOO) and introduces a novel concept of the Utopian front, an extension of Pareto fronts, to analyze the conservation and diversification of the BMP mechanism between *Drosophila* and zebrafish. By applying Pareto front analysis, we explore how the biophysical properties of the BMP pathway have been conserved in *Drosophila* and zebrafish systems.

We identified minimal biophysical requirements that reconcile known geometric and temporal differences between *Drosophila* and zebrafish while producing experimental observations. Through the Pareto analysis, we identified a set of biophysical parameters that generated the best compromise to fit both *Drosophila* and zebrafish BMP systems.

Experimental data further suggest a divergence between the *Drosophila* and zebrafish BMP pathways. The novel concept of Utopian fronts provides a new framework to (i) assess system constraints in fitting different objectives and (ii) distinguish between variant and invariant components within a system. By combining Pareto optimization and Utopian analysis, we systematically varied a subset of parameters (variant component) while keeping others constant at their Pareto optimal values (invariant). Geometric differences alone proved insufficient to explain the evolutionary divergence of BMP mechanisms. Instead, a balance between geometry and biophysical properties was required to produce BMP patterns in *Drosophila* and zebrafish. Utopian analysis identified the characteristic length of the BMP-Sog and the Sog cleavage rate as key drivers of BMP mechanism divergence. These parameters, previously shown to be important in BMP shuttling in *Drosophila*, help generate the steep and sharp BMP gradient that is evidently different from the zebrafish BMP mechanism. The evolutionary changes, especially in BMP-specific parameters, in addition to differences in geometry and complex incorporation of positive feedback (in *Drosophila*) and Noggin regulation (in zebrafish), contribute to species-specific BMP mechanisms. Overall, our findings demonstrate how Pareto optimization and Utopian analysis provide a structured approach to studying complex systems. This framework not only helps investigate evolutionary system dynamics but also identifies components of systems selected for conservation and those selected for divergence. More broadly, this methodology could be extended to other biological systems where mechanisms can be generalized through a common model or common parameter.

## ACKNOWLEDGEMENT

This work is based upon efforts supported by the EMBRIO Institute, contract 2120200, a National Science Foundation (NSF) Biology Integration Institute. This research was supported in part by the NSF grant 2422229 awarded to L.L and D.M.U.

## Notes

### Competing Interest Statement

The authors have declared no competing interest.

