## Supplemental figures for "Digital Cousins: Simultaneous Optimization of One Model for BMP Signaling in Distant Relatives Reveals Essential Core"

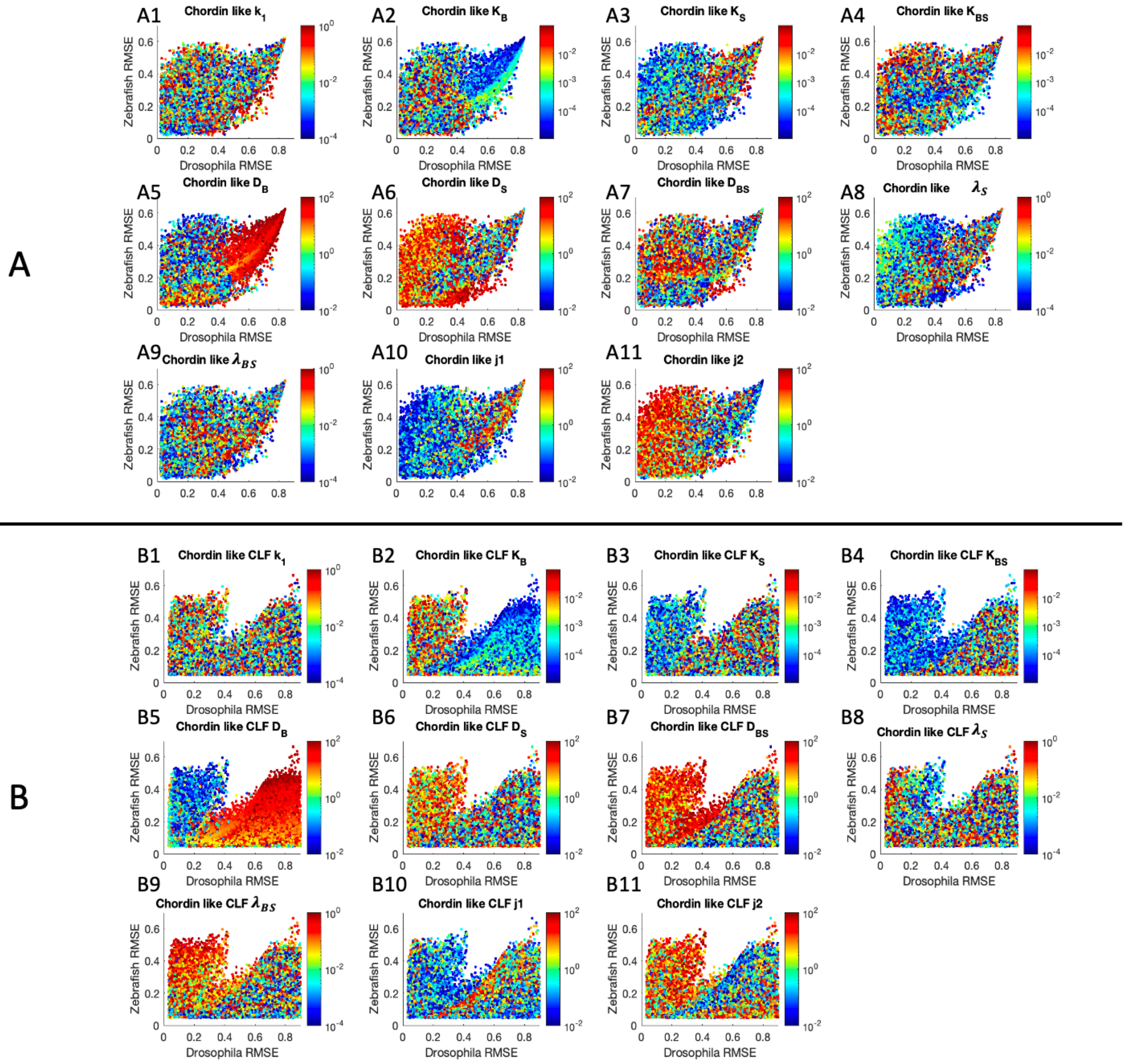


*Figure S1 Parameter trend analysis for all the simulations results against Chordin like and Chordin like Chd(-/-) fitting. Figure group A shows the parameter tendency in Utopian space against Chordin like fitting for 11 independent parameters, including A1, $k_1$( forward reaction rates for Bmp and Chd/Sog); A2, $K_B$ (Decay rate for BMP); A3, $K_S$ (Decay rate for Chd/Sog); A4, $K_{BS}$ (Decay rate for BMP-Chd/Sog complex); A5, $D_B$ (Diffusion rate for BMP); A6, $D_S$ (Diffusion rate for Chd/Sog); A7, $D_{BS}$ (Diffusion rate for BMP-Chd/Sog complex); A8, $\lambda_S$ (Tld processing rate of Chd/Sog); A9, $\lambda_{BS}$ (Tld processing rate of BMP-Chd/Sog complex); A10, J1(Production rate of Bmp); A11, J2 (Production rate of Chd/Sog). Group B shows the parameter tendency in Utopian space against Chordin like data with Chd/Sog(-/-) fitting for 11 independent parameters, respectively. Notably, the RMES range is limited to [0,0.6] for Zebrafish fitting and [0,0.8] for Drosophila fitting for better visualization.*

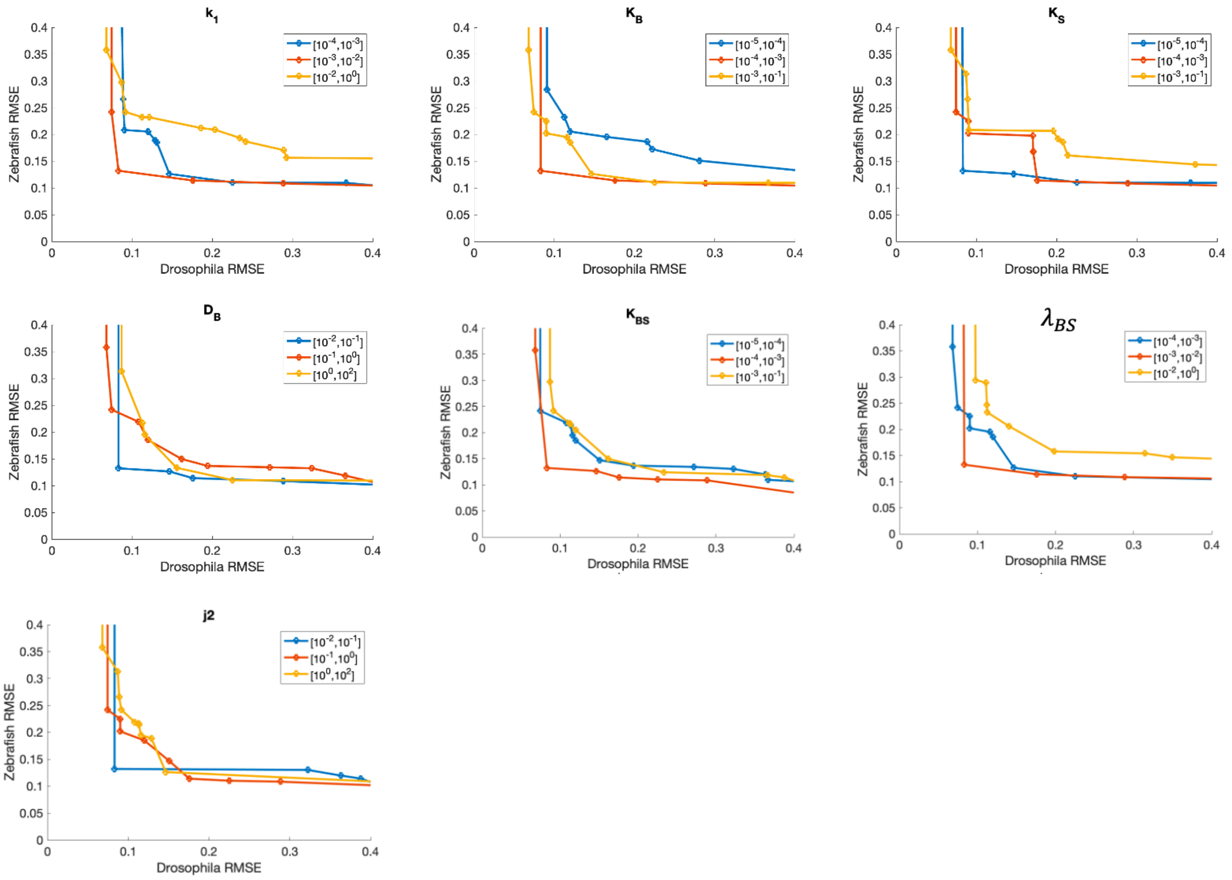


*Figure S2 Utopian front of WT + Sog/Chd(-/-) fit for the individual parameters in specific ranges. $k_1$( forward reaction rates for Bmp and Chd/Sog); $K_B$ (Decay rate for BMP); $K_S$ (Decay rate for Chd/Sog); $K_{BS}$ (Decay rate for BMP-Chd/Sog complex); $D_B$ (Diffusion rate for BMP); $\lambda_S$ (Tld processing rate of Chd/Sog, J2 (Production rate of Chd/Sog).*
